# Structure and epitope mapping of the conformational anti tau antibody DC11

**DOI:** 10.64898/2026.09.01.748351

**Authors:** Stefana Njemoga, Leon Jenner, Viliam Volko, Adam Polak, Lubica Fialova, Tomas Augustin, Jozef Hanes, Aneta Kozelekova, Radek Crha, Lucia Il’kovičková, Rostislav Skrabana, Pavel Kaderavek, Petr Kolenko, Tomas Smolek, Branislav Kovacech, Jozef Hritz, Ondrej Cehlár

**Author notes:** These authors contributed equally (Stefana Njemoga, Leon Jenner).

## Abstract

Conformational antibody DC11 was previously shown to discriminate between physiological full length tau proteins and misfolded truncated tau proteins. It was also shown to catalyze in vitro tau aggregation, suggesting a connection with the pre-aggregation conformation of tau proteins. We have crystallized the Fab fragment of the DC11 antibody and characterized its binding with truncated tau proteins using ELISA, NMR and crosslinking mass spectrometry. The presumed model of the complex of DC11 antibody and truncated tau protein was obtained by docking tau321-391 conformations from coarse grained MD simulation into the antibody paratope.

## 1. INTRODUCTION

Tau is an intrinsically disordered protein that regulates the dynamics of microtubules, encoded by the microtubule associated protein tau (MAPT) gene. In neurodegenerative tauopathy, tau misfolds and deposits in the brain tissue in the form of amyloid filaments, where each tauopathy is characterized by a specific tau amyloid fold. Alternative splicing of the MAPT gene gives rise to six CNS tau isoforms, that have 3 or 4 microtubule binding repeats (3R or 4R respectively). Tauopathies are than characterized by the presence of deposits composed of either 4R tau proteins (Corticobasal degeneration) or 3R tau proteins (Picks disease), or there are mixed deposits composed of both 3R and 4R tau proteins (Alzheimer’s disease (AD), Chronic traumatic encephalopathy). [1]

Pathological conformations of tau range from misfolded monomeric states to soluble oligomeric species and high-Mw tau fibrils, the final product of tau aggregation, deposited as neurofibrillary tangles in the diseased brain. Transient pathological structures sampled during the conversion from healthy, disordered tau conformations through aggregation-prone conformations to fibrils can be selectively captured by specific monoclonal antibodies (mAbs). Such mAbs, recognizing epitopes specific to misfolded tau monomers or oligomers, are of great importance because they distinguish healthy from toxic tau species and enable the targeting of tau aggregation at its earliest stages. The potential of monoclonal antibodies is not only therapeutic; disease-specific antibodies can also reveal disease-relevant tau epitopes, their misfolding mechanisms, and structures, which can be further used to design targeted anti-tau therapies.

Specific monoclonal antibodies developed against AD-tau species, such as antibody MN423, have revealed that the structured core of tau paired helical filaments (PHF) is composed of truncated tau proteins [2,3]. One of these truncated fragments, termed dGAE (tau297-391), has recently been widely used because of its ability to attain the amyloid fold of AD PHF in vitro [4]. Antibody MN423 is specific for the E391 truncated tau species and was previously crystallized in complex with the tau hexapeptide ^386^TDHGAE^391^ [5].

The conformational antibody DC11 was generated in mice immunized with hippocampal extracts from AD brain and specifically recognizes truncated tau variants (tau151-421 – minimal truncation) that are highly aggregation-prone and sufficient to initiate tau pathology in a rat tauopathy model [6]. Taken together, the high specificity for pathologically modified tau, the requirement for truncation, and the catalytic pro-aggregatory effect on tau suggest that the DC11-reactive conformation represents a transient pre-aggregation state between physiologically disordered and aggregation-competent tau [7]. In addition, both DC11 and MN423 antibodies were shown to promote in vitro tau amyloid formation in aggregation assays with truncated tau [8].

## 2. METHODS

### 2.1 Production of DC11 Fab in CHO cells

CHO cells were co-transfected with pCMV_3′UTR plasmids encoding the heavy and light chains of Fab DC11 and MN423 at a 1:1 ratio using polyethyleneimine (PEI). The final DNA concentration was 4 μg/mL, and PEI was added at 8 μg/mL, corresponding to the optimized PEI:DNA ratio of 2:1, as previously described [9]. CHO cells were cultivated in BalanCD Growth A medium supplemented with 6 mM Glutamine in a 37 °C humidified environment with 8% supplemented CO_2_ (Irvine Scientific). The cells were transferred into BalanCD Transfectory medium supplemented with 8 mM glutamine 24 h prior to transfection (Irvine Scientific). The cells were transferred into a 32 °C humidified environment with 5% CO_2_ 24 h after post-transfection. Secreted DC11 Fab was purified from culture supernatant by affinity chromatography using a 5 mL HiTrap Protein G column (GE Healthcare). For MN423 Fab, a 1 mL HiTrap Protein L column was used. The pH of the supernatants was adjusted to 8.0 by gradual addition of 1 M Tris-HCl (pH 9.0). The samples were subsequently centrifuged at 21 000 × g for 20 min and filtered through a 0.2 μm membrane filter prior to chromatography. The clarified supernatant was applied to the corresponding columns. After washing with 1× PBS to remove unbound proteins, Fabs were eluted using 100 mM glycine-HCl (pH 2.7). Eluted fractions were immediately neutralized by the addition of 1/10 volume of 1 M Tris-HCl (pH 9.0). Desalting and buffer exchange were performed using a HiPrep 26/10 desalting column (GE Healthcare) into 10mM Tris-HCl pH 7.2, 50 mM NaCl solution. Fabs were further concentrated using Amicon 10K Ultra centrifugal filters (Sigma-Aldrich) and the final concentration was calculated from absorbance at 280 nm.

### 2.2 Expression and purification of truncated tau proteins

Truncated tau variants dGAE (Tau297–391) and Tau321–391 and dGAE deletion mutants and C322S mutants were expressed in *E. coli* strain BL21(DE3) (Sigma-Aldrich) from a pET-17 expression vector and purified from bacterial lysates by a series of ion exchange and size exclusion chromatography steps according to published protocol [10]. Purified tau proteins were stored in PBS pH 7.4 at −80 °C under an argon atmosphere to prevent disulfide formation. The concentration of purified tau proteins was determined from the absorbance at 205 nm using a DU 640 spectrophotometer (Beckman Colter, USA). Tau proteins and their respective PBS blank were diluted in 10 mM phosphate buffer, pH 7.2, with 0.0025% Tween 20. The absorption coefficient was calculated using the Protein Parameter Calculator webserver [11]. The purity and identity of purified tau proteins were verified by sodium dodecyl sulfate– polyacrylamide gel electrophoresis (SDS-PAGE) and by mass spectrometry (MS) analysis.

#### 2.2.1 ^13^C, ^15^N isotope labelling of tau proteins

For isotope labelling, cells were cultured in M9 minimal medium supplemented with 2 g/L ^13^C glucose and 1 g/L ^15^N ammonium chloride (Cambridge Isotope Laboratories, USA) using the Marley protocol [12], where the culture was initially grown in LB medium and concentrated 4-fold into the minimal medium before induction with IPTG.

### 2.3 Crystallization

DC11 Fab crystals were grown in 0.1 M MIB (malonic acid : imidazole : boric acid in 2:3:3 molar ratio), pH 6.0 and 25% w/v PEG 1500, formulated as the condition B3 of PACT Premier screen HT (Molecular Dimensions, UK) with a final protein concentration of 8 mg/mL. Crystallization experiments were performed manually according to our previously published protocol [13] using the sitting-drop vapor diffusion method in 96-well, two-drop MRC crystallization plates (Molecular Dimensions). 80 μL of reservoir solution was dispensed into the reservoir wells. Drops of 0.5 μL reservoir solution were transferred into the crystallization wells using a motorized eight-channel pipette (Gilson). Subsequently, 0.5 μL of protein solution was added to each drop. Plates were sealed with adhesive film (Greiner Bio-One) and kept in a temperature-controlled room at 22 °C (295 K). Crystal growth was monitored periodically using a Nikon SMZ645 microscope.

#### 2.3.1 Diffraction data collection, processing, and structure determination

Crystals were harvested using nylon cryoloops (Hampton Research) or polymer Micro Loops (Mitegen), briefly immersed in 10% trehalose, then in Paratone-N cryoprotectant, and flash-cooled in liquid nitrogen. X-ray data were collected at beamline P13 (EMBL) at the PETRA III synchrotron facility (DESY, Hamburg) using an X-ray wavelength of 0.9763 Å, and an EIGER X 16M detector (Dectris, Switzerland). After initial characterization using 4 scans with 90° rotation, the full datasets were measured, usually containing 3600 images with 0.1° rotation (fine slicing). Collected X-ray diffraction images were processed using the automated diffraction data-processing pipeline autoPROC at the beamline. Diffraction data indexing and integration were performed with XDS. The most probable space group was determined using POINTLESS. Data scaling and merging were performed with AIMLESS, and diffraction anisotropy was analyzed with STARANISO. The structure was solved by molecular replacement using PHASER with [PDB ID: 7T62] as the search template. Iterative model building and manual corrections were performed in Coot, followed by structure refinement using REFMAC. Free R value was used as a validation metric. The raw data images were deposited into the zenodo repository [14] within the Macromolecular crystallography community.

### 2.4 ELISA

#### 2.4.1 Sandwich ELISA

300 ng of purified DC11 antibody per well was dispensed onto a microtitre ELISA plate in 100 μl of PBS. The plate was incubated at 25 °C for at least 2 hours or overnight at 4 °C. The plate was blocked with 10 mM Tris buffer pH 8.8 with 0.05% Tween 20 (Tris-T) for 30 minutes at 25 °C. In a polypropylene plate, a two-fold serial dilution of tau protein in 1× Tris-T was prepared, ensuring that at least 120 μl of each concentration was prepared. The highest (first) concentration in the tau protein dilution series was 500 nM. After blocking, the Tris-T buffer was removed and 100 μl of tau protein was transferred from each dilution to the antibody-coated plate. In the next step, the ELISA plate was incubated for 2 hours at 25 °C on a rotary platform set to 230 rpm. The plate was washed three times with 1× Tris-T (using an automatic washer, Athos Fluido, set to 500 μl of wash buffer per wash). Then 100 μl per well of the HRP-conjugated secondary antibody DC25 (Axon Neuroscience) diluted 1:5000 in Tris-T was added, and the plate was incubated for 1 hour at 25 °C on a rotary platform set to 230 rpm. Li-COR ELISA HRP substrate was prepared 5–10 minutes before use according to the manufacturer’s instructions. The plates were washed three times with Tris-T, and 100 μl per well of substrate was added and plates were incubated at room temperature for 10–20 minutes in the dark. The reaction was stopped by adding 20 μl per well of 2M H□SO□ and the absorbance at 700 nm was subsequently measured (Odyssey infrared imaging system). EC□ □ values were determined based on the immunoreactivity using GraphPad Prism version 3.02 for Windows.

#### 2.4.2 Competitive ELISA

100 ng per well of purified tau dGAE (tau297–391) was dispensed in 100 μl of PBS onto a microtitre ELISA plate. The plate containing tau protein was incubated for at least 2 hours at 25 °C or overnight at 4 °C. The plate was then blocked with 1× Tris-T for 30 minutes at 25 °C. A two-fold serial dilution of tau protein deletion mutants (competitor) in Tris-T buffer was prepared in a polypropylene plate. The highest concentration in the competitor dilution series was 20 μM. 100 ng per well of the DC11 antibody in 100 μl of Tris buffer was added to the diluted competitor. After blocking, 100 μl per well of each dilution of the competitor–antibody mixture was transferred onto the tau297–391 coated plate. The plate was then incubated for 2 hours at 25 °C on a rotary platform at 230 rpm. After washing with Tris-T, 100 μl per well of anti-mouse antibody conjugated with HRP was added, diluted 1:5000 in Tris-T, and incubated for 1 hour at 25 °C on a rotary platform at 230 rpm. Further steps were the same as in 2.4.1.

### 2.5 NMR experiments

#### 2.5.1 NMR assignment of dGAE

NMR chemical shift assignment of dGAE (prepared as in [15]) was performed using 3D and 5D nonuniformly sampled (NUS) NMR experiments [16,17]. The 3D and 5D CACONCACO and HC(CCTOCSY)CACON spectra were acquired for 1 mM [^13^C, ^15^N] dGAE in 50 mM sodium phosphate pH 6.9, 5 mM TCEP and 10% D_2_O at 20 °C using a 700 MHz NMR spectrometer Bruker Avance NEO equipped with TXO 5 mm triple-resonance ^1^H/^13^C/^15^N cryoprobe optimized for ^13^C detection. The NMR assignment is deposited under BMRB entry 53992.

#### 2.5.2 NMR titration experiments

The binding epitope between dGAE and DC11 Fab was mapped using 2D HSQC and 3D HNCO (with NUS) experiments measured using a 950 MHz NMR spectrometer Bruker Avance NEO with 5 mm TCI triple-resonance ^1^H/^13^C/^15^N inverse cryoprobe for ^1^H detection, under the control of Topspin 4.1 software (Bruker). The 2D HSQC spectra were obtained using the standard Bruker pulse sequence hsqcetf3gp [18] and the 3D HNCO spectra were acquired using the hncogpwg3d_t2sc pulse sequence [19–21], featuring watergate water suppression [22], gradient filters, and semiconstant time ^15^N evolution. NMR spectra for samples containing DC11 Fab and [^13^C, ^15^N] dGAE in molar ratios 0:1 (0:590 μM), 1:1 (250:250 μM), 2:1 (320:160 μM) and 4:1 (320:80 μM) were acquired at 5 °C in PBS pH 7.4, 10% D_2_O in the presence of a commercial protease inhibitor cocktail (Roche) to prevent protein degradation. Data were processed using NMRpipe [23]. The NUS dimensions were processed using the SMILE package [24]. NMRFAMSPARKY software [25] was used to fit and integrate 3D HNCO peak volumes. Dilution effects were accounted for and peak volume ratios (between dGAE in bound and free state) were calculated using Microsoft Excel and QTIplot (IonDev software). The NMR data are deposited as BMRB entry 53993.

Similar experiments were performed with MN423 Fab and [^13^C, ^15^N] dGAE in molar ratios 0:1 (0:240 μM) and 1:1 (240:240 μM) in PBS pH 7.4, 10% D_2_O at 25 °C as a control experiment.

### 2.6 Crosslinking mass spectroscopy

Crosslinking mass spectroscopy (XL-MS) experiments were used to identify residues in dGAE within close proximity to DC11 Fab in complexes (Cα-Cα distance between the cross-linked residues of approx. 27 Å). These experiments rely on reactivity between MS-cleavable crosslinker disuccinimidyl dibutiric urea (DSBU) and K, S, T and Y sidechains or the N-terminal amine of any amino acid. XL-MS experiments were performed according to the protocol used in reference [26]. Briefly, 0:1, 1:0, 1:2, 2:1 and 1:1 samples of DC11 Fab and non-labelled dGAE were incubated with 200 μM DSBU for 30 min at 37 °C, from which 1:2 samples were chosen for final analysis. The reaction was quenched with 1 mM Tris-HCl, pH 8.0 for 10 min, then resolved on 12.5% SDS-PAGE gel (200 V, 60 min). Bands consistent with the mass of both DC11 Fab chains plus dGAE after Coomassie brilliant blue staining were excised, digested with trypsin and then analysed by LC-MS/MS using a RSLCnano system, connected to an Orbitrap Fusion Lumos Tribrid mass spectrometer (Thermo Fisher Scientific). MS mass/charge spectra were converted to Mascot generic files (.mgf) using ProteoWizard [27] for subsequent cross-link analysis by MeroX [28].

### 2.7 Antibody-Tau docking

The crystal structure of DC11 Fab (PDB ID: 9H8H) determined by X-ray crystallography was used to prepare a scFv model used as the receptor (target) structure for docking using the HADDOCK web server [29]. Variable domains of heavy and light chain were connected with a GGGSGGGSGGS linker (an offset of 123 has to be used to get light chain residue numbers consistent with numbering of the DC11 Fab model). Representative conformers of Tau321–391 obtained from coarse-grained MD simulations [30] were backmapped using sirah_backmap implemented in AmberTools25 and used as ligand structures. Tau residues 371–386 identified by NMR epitope mapping were designated active residues, together with the complementarity-determining regions (CDRs) of the DC11 Fab, which were assigned using the AbRSA platform [31] based on the Chothia scheme. All remaining docking parameters were kept at their default values in HADDOCK 2.4.

## 3. RESULTS

### 3.1 Crystal structure of DC11 Fab fragment

High-throughput screening of crystallization conditions was performed in a sitting-drop format using multiple commercially available crystallization screens. We obtained multiple crystallization hits for DC11 Fab. The structure was solved from the crystal grown during the initial screening in the condition from PACT premier crystallization screen, without the need for further optimization. The statistics of the processed diffraction data are shown in the Table 1.

**Table 1.** Diffraction data collection statistics of DC11 Fab.

|  |  |
| --- | --- |
| *Diffraction source* | PETRA III/P13 (MX1) |
| *Wavelength (Å)* | 0.97626 |
| *Temperature (K)* | 100 |
| *Detector* | DECTRIS EIGER X 16M |
| *Space group* | $P1\ 2_1\ 1$ |
| *Length $a,b,c$ (Å)* | 40.901 88.445 57.464 |
| *Angle $\alpha,\beta,\gamma$ (°)* | 90 94.803 90 |
| *Resolution (Å)* | 48.07–1.33 (1.43–1.33) |
| *Total reflections* | 417959 |
| *Unique reflections* | 70143 (3508) |
| *R<sub>merge</sub> (%)* | 0.092 (1.088) |
| *(I/$\sigma$(I))* | 9 (1.4) |
| *Completeness (%) - ellipsoidal* | 90.2 (53.9) |
| *Multiplicity* | 6.0 (5.9) |
| *Mosaicity (°)* | 0.2 |
| *Overall B factor (Å<sup>2</sup>)* | 21.224 |
| *R<sub>meas</sub> (%)* | 0.102 (1.192) |
| *Matthew's coefficient (Å<sup>3</sup>/Dalton)* | 2.19 |
| *Solvent content (%)* | 43.91 |

The structure of DC11 Fab was refined to final R_work_ and R_free_ values of 0.145 and 0.196, respectively. The refined model exhibited a mean isotropic B-factor of 21.2 Å^2^, consistent with a well-ordered crystal structure. The final structure was deposited into the Protein Data Bank under PDB accession code 9H8H (Fig. 1). From the surface electrostatics representation, we can distinguish the electropositive bottom of the antibody binding site surrounded by the electronegative wall formed by the heavy and light chain CDR loops.

**Figure 1.**
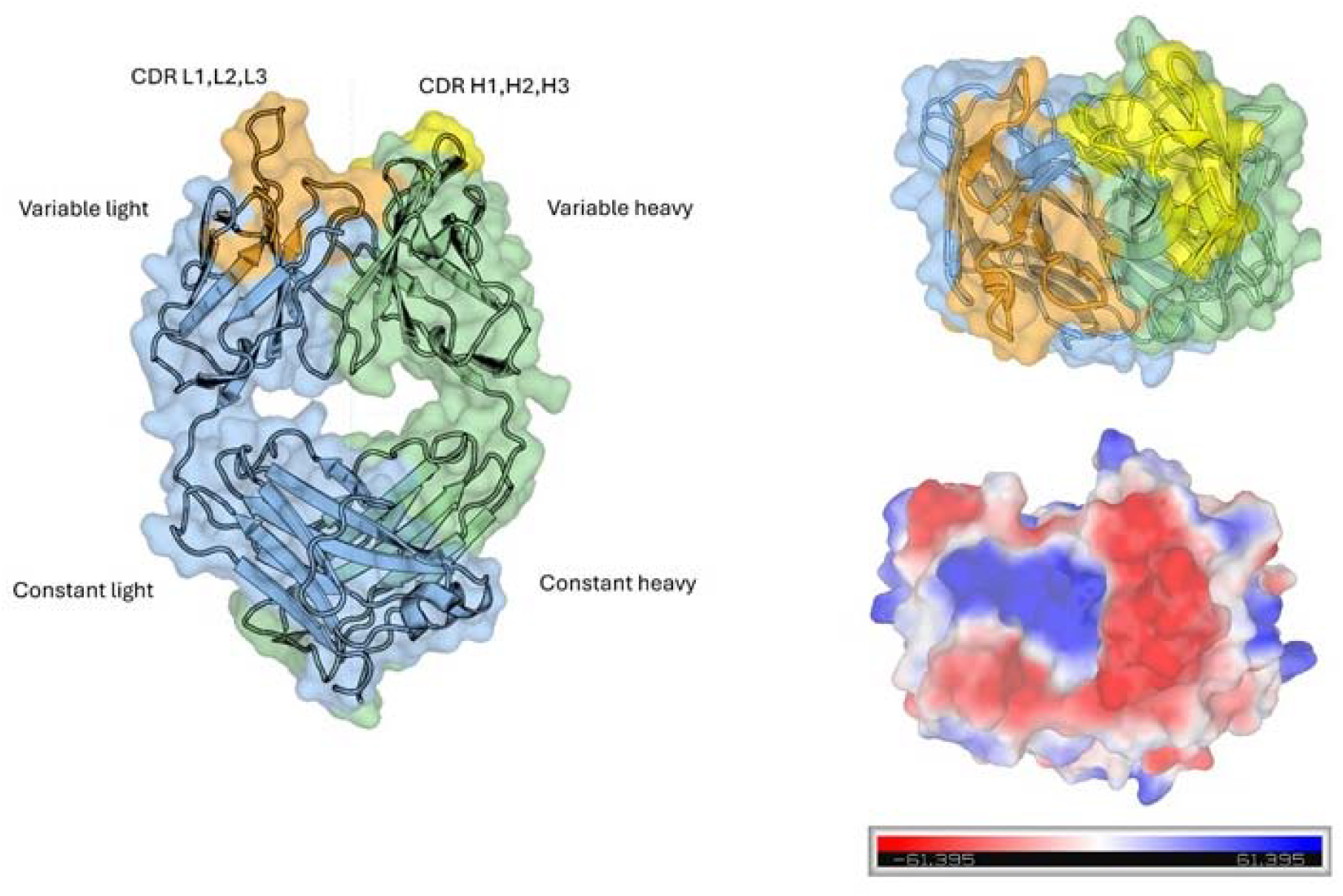
Crystal structure of the DC11 Fab fragment. The structure consists of light and heavy chains (colored blue and green), each with constant and variable domains. The three complementarity-determining region (CDR) loops of both chains are highlighted in orange and yellow. On the right there is 90° rotated image viewing from top into the paratope together with surface vacuum electrostatics representation (bottom right).

The AbRSA web tool was used to identify DC11 complementarity-determining regions (CDRs) using the Chothia and IGMT algorithms. These predicted that the DC11 antibody has a relatively short CDRH3 loop composed of only 5 amino acids (according to Chothia, Fig 2).

**Fig 2.**
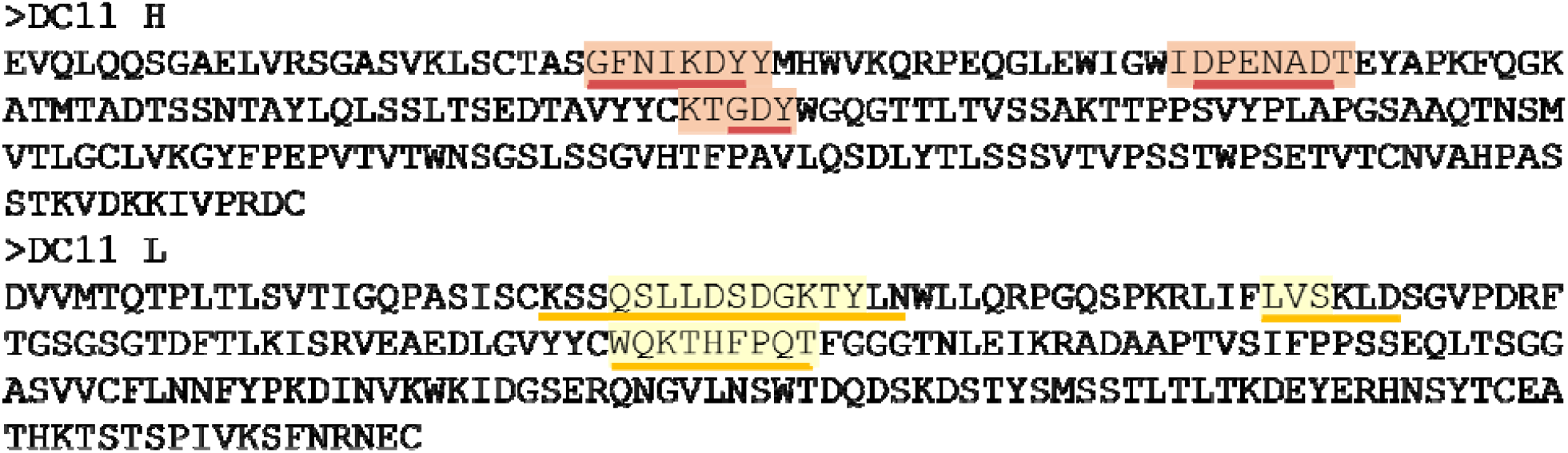
Complementarity-determining region (CDR) sequences of the DC11 Fab. identified according to the IMGT and Chothia numbering schemes. IMGT-defined CDRs are indicated by orange (heavy chain) and yellow (light chain) boxes, whereas Chothia-defined CDRs are marked by underlines.

### 3.2 DC11 Fab epitope mapping by ELISA

#### 3.2.1 Sandwich ELISA

Sandwich ELISAs were performed to confirm the preference of DC11 antibody for truncated tau proteins over full length tau proteins. Antibody DC11 was immobilized in the microplate wells. The binding of tau proteins was detected using the HRP labelled antibody DC25 (Axon Neuroscience, Slovakia). The results of this ELISA assay showed a clear preference of DC11 for truncated tau proteins, dGAE and longer truncated tau proteins tau151-391/4R [32] and tau151-391/3R [33] over the full length 4R tau 2N4R (Fig. 3).

**Fig 3.**
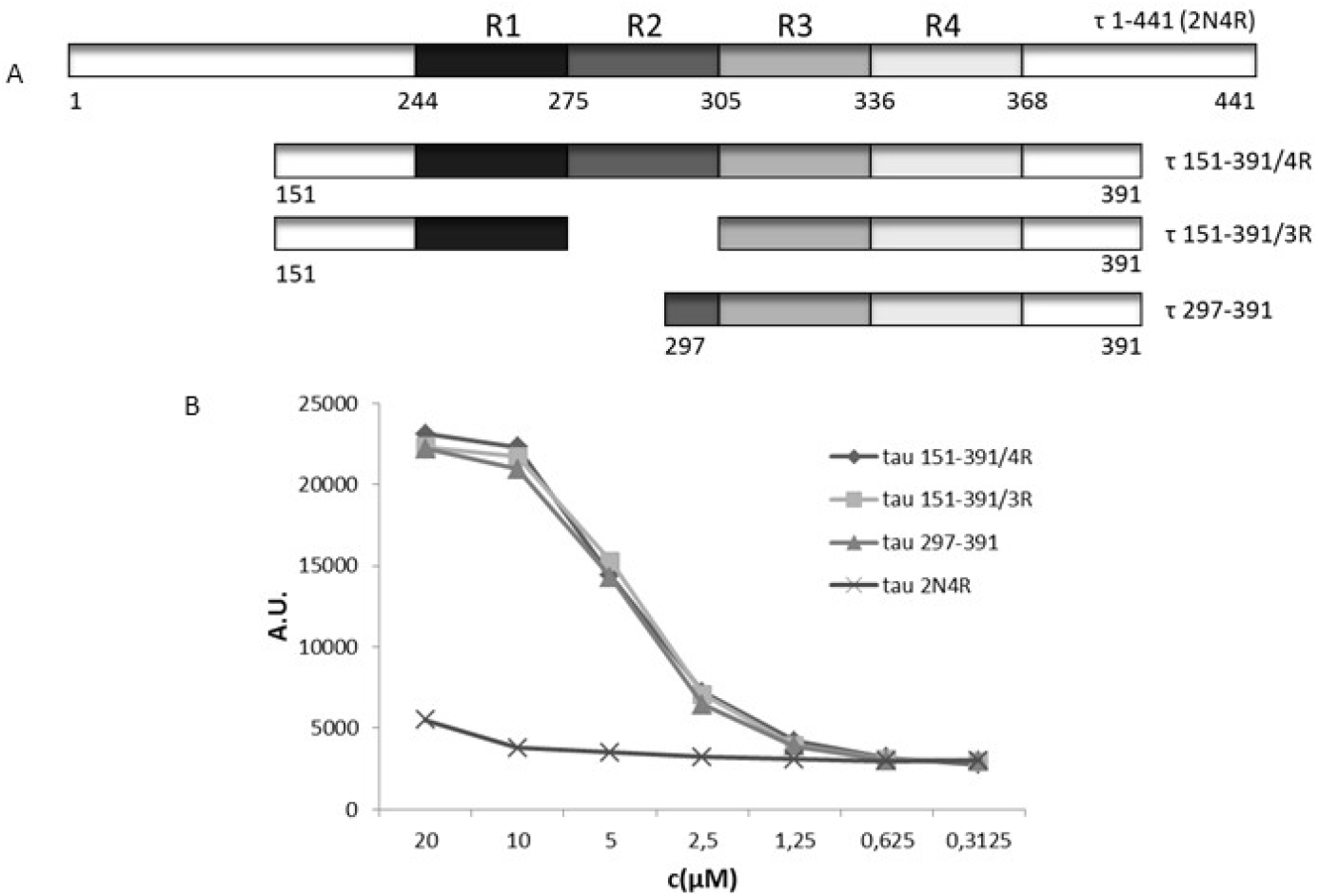
DC11 recognizes preferentially tau proteins that are truncated at both the N-and C-terminal ends. A) Diagram of tau proteins used for immunoreactivity of DC11 in sandwich ELISAs. The numbering of amino acids is that of full-length human tau τ1-441 (2N4R). (B) Graph shows interaction of DC11 antibody with double truncated tau proteins and full-length tau τ1-441 (2N4R). Individual curves reflect immunoreactivities of respective tau proteins.

#### 3.2.2 Competitive ELISA

A panel of dGAE deletion mutants was prepared to further characterize the DC11 tau epitope using the competitive ELISA method, where the reactivity of mutants was compared to the reactivity of tau dGAE, immobilized on microplates. EC_50_ values were derived from competition curves. The reactivity of N-terminally truncated variants of dGAE (starting at tau residues 306, 316, 321 and 326 [30]) confirmed previous western blot results [6], where the antibody also bound all but the shortest tau326-391 fragment (Fig. 4).

**Fig 4.**
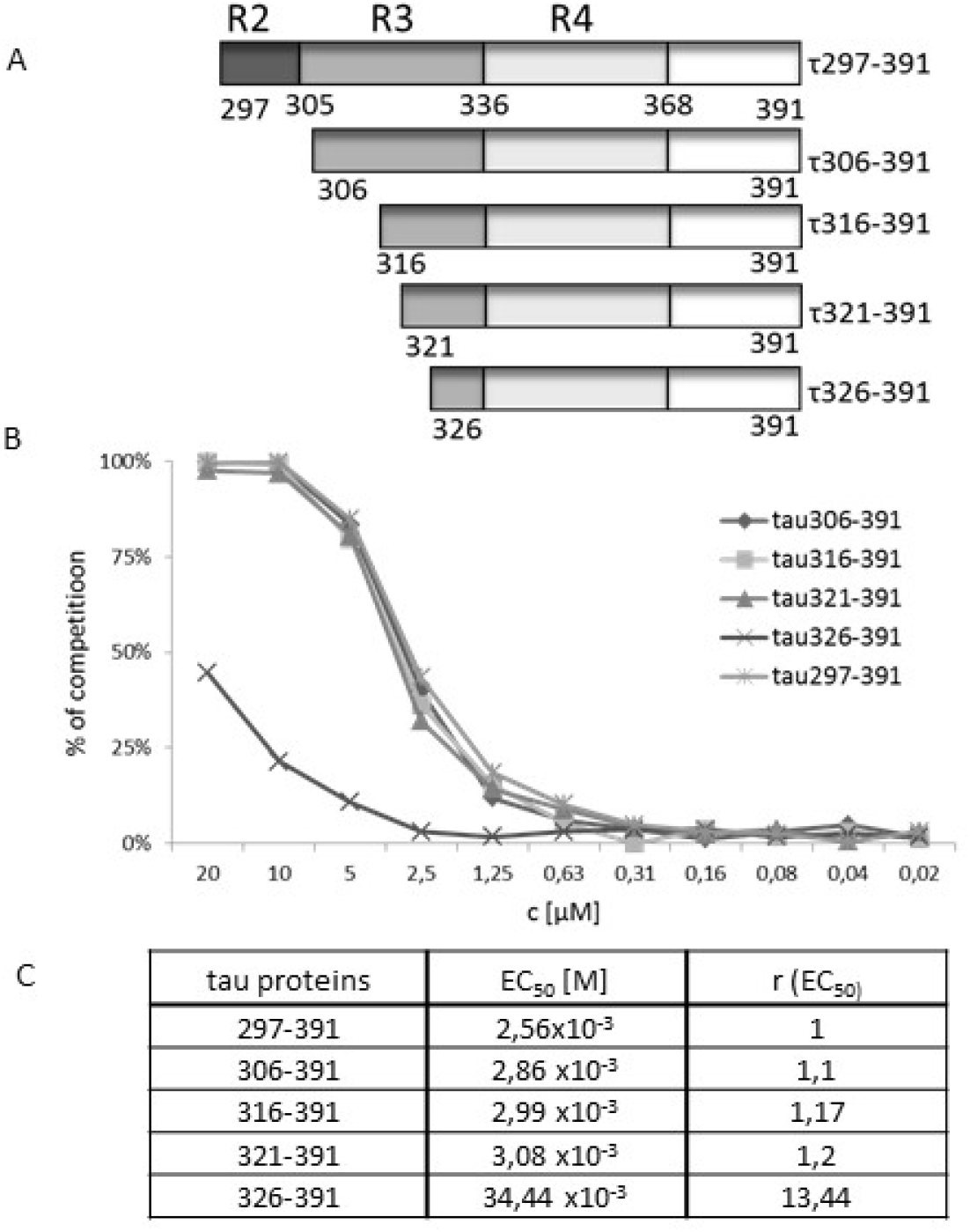
Identification of sequences contributing to the epitope of DC11 located in the third tau repeat. A) Structural diagram of N-terminal deletion mutants derived from τ297-391 used for immunoreactivity of DC11 in competitive ELISA. B) Graph shows competition of N-terminal deletion mutants of τ297-391 for binding with DC11. Individual competition curves reflect immunoreactivities of DC11 with respective mutants. C) EC_50_ values were calculated for measure the affinity of DC11; ratio between EC_50_ of respective deletion mutant and EC_50_ of τ297-391 (r) indicates decrease in relative affinity of mutant (rfold) in relation to full affinity of τ297-391.

The sequence 321-325, which we have previously shown to be important for the aggregation of truncated tau proteins [30], seems to be also crucial for the binding of tau to the DC11 antibody. Therefore, a set of deletion mutants was prepared to test the contribution of short peptide segments to the molecular interface between tau and the DC11 antibody (Fig. 5). The results showed substantially lowered reactivity upon deletion of either the 321-325 segement or 321-325 and 306-311 segments (Fig. 5). This result is consistent with the idea that the 321-325 sequence forms part of the tau conformational epitope recognized by the DC11 antibody.

**Fig 5.**
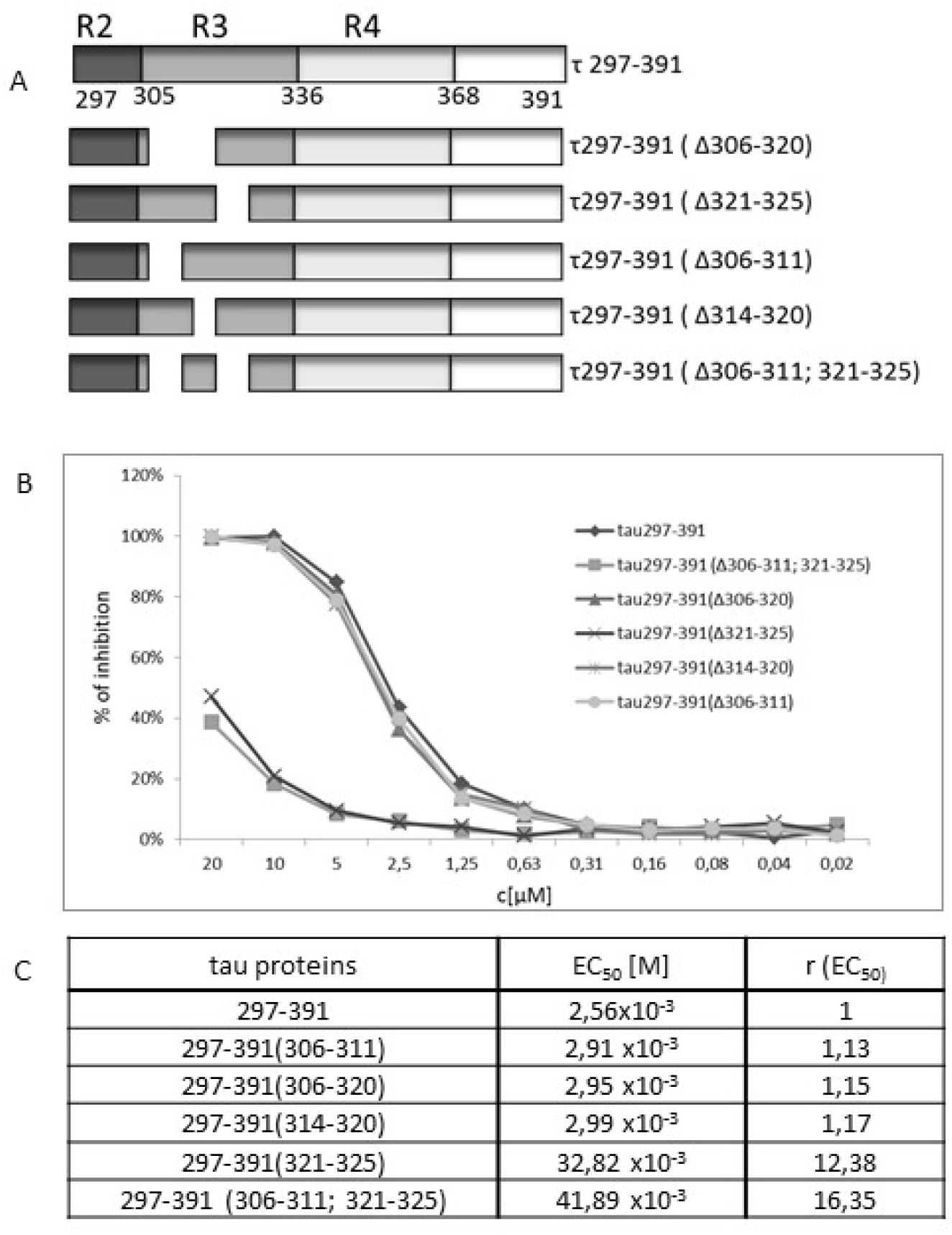
Identification of sequences contributing to the epitope of DC11 antibody located in the third tau microtubule binding repeat. A) Structural diagram of internal deletion mutants derived from τ297-391 used for immunoreactivity of DC11 in competitive ELISAs. B) Graph shows competition of internal deletion mutants of τ297-391 for binding with DC11. Individual competition curves indicate DC11 immunoreactivities with respective mutants. C) EC_50_ values were calculated to measure the affinity of DC11; the ratio between EC_50_ of respective deletion mutants and EC_50_ of τ297-391 (r) indicates decrease in relative affinity of mutant (r-fold) in relation to full affinity of τ297-391.

One cystein residue (C322) is located within the identified 321-325 region. Several aggregation mechanisms of tau have already been reported, including the formation of intermolecular disulfide bonds, which lead to different morphologies of the resulting tau fibrils. To test whether the C322 plays a role in DC11 recognition, Tau321-391 and dGAE with the C322S point mutation have been prepared and have shown no significant change in binding affinities (Fig. S1).

C-terminal deletions of dGAE were also probed using tau variants tau297-384 and tau297-376, where the shorter tau fragment showed lowered reactivity with DC11 (Fig. S2).

### 3.3 DC11 Fab epitope mapping by NMR and XL-MS

The binding interface between DC11 Fab and tau dGAE was mapped using NMR spectroscopy. As IDPs are known to feature considerable signal overlap, known assignments for dGAE [34] were confirmed, and missing signals assigned (Fig. S3) using the IDP-specialised 5D assignment experiments previously used for full-length tau [26]. The 3D HNCO spectrum of isotopically enriched [^13^C,^15^N] protein shows signals, each of which can be assigned to a single backbone amide proton-nitrogen pair, correlated to the backbone carbonyl group of the preceding amino acid. As DC11 Fab is titrated into the sample in a 1:1 to 4:1 molar ratio, some of the signals are attenuated (Fig. 6A) which could indicate binding or other changes of dGAE mobility, similar to those observed in [26,35] for full length tau and a protein binding partner. Signal areas decrease in a DC11-dependent manner for most residues aa369-391 in the C-terminus of dGAE, suggesting this as an extended binding surface. Consistent with this, parallel crosslinking mass spectrometry (XL-MS) experiments with non-labelled dGAE and DC11 Fab, mixed in a 2:1 ratio, identified crosslinks indicating DC11 proximity across the dGAE sequence, with more unique crosslinks towards the C-terminus (Fig. 6B, S4). Residues aa322-326 show a similar loss in NMR signal, consistent with ELISA data that show these residues as necessary for high affinity binding. Hydrophobic residues V300, I308, V313 and V339 also have lower intensity in the presence of DC11, suggesting some interaction. Similar loss of signal, albeit for different residues, was observed for [^13^C,^15^N] dGAE in a 1:1 complex with MN423 FAB (Fig. 6C). 3D HNCO data in this case indicate a primary interaction site at aa385-391, consistent with the binding of aa387-391 to MN423 Fab observed in an X-ray diffraction derived structural model [5] (PDB 2V17).

**Figure 6.**
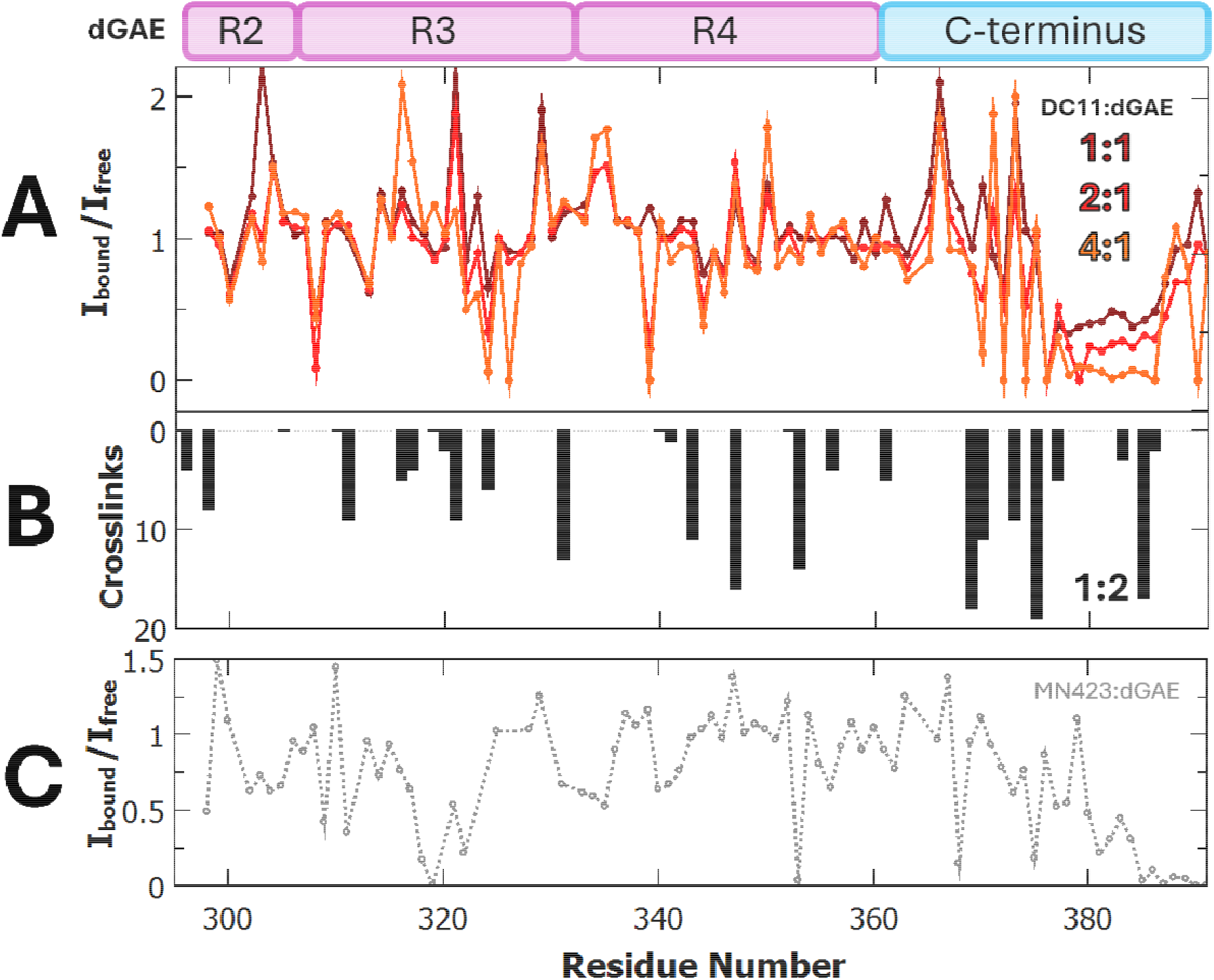
– Binding interface of the dGAE:DC11 complex. Binding interface between tau dGAE and DC11 FAB investigated by **A)** NMR titration or **B)** XL-MS. **A)** [^13^C,^15^N] dGAE signal areas (I_free_) in HNCO experiments were divided by signal volumes in the presence of varying ratios of DC11 FAB (I_bound_) as annotated. Loss of NMR signal volume is observed for residues that interact with DC11 Fab. **B)** XL-MS data of 1:2 DC11 FAB and unenriched dGAE, showing counts for the number of unique crosslinks detected per amino acid (black bars), with grey dashed line indicating residues chemically incapable of crosslinking. **C)** Comparative NMR binding interface as A) for 1:1 [^13^C,^15^N] dGAE and MN423 FAB (grey dashed lines).

### 3.4 Antibody-tau docking

Based on the obtained structural data, we performed antibody–tau docking using the HADDOCK web server, a multi-stage protocol that includes rigid-body energy minimization, semi-flexible refinement, and final refinement in explicit solvent. In the initial stage, the interacting molecules are randomly oriented and translated to sample possible docking poses. These are subsequently refined to optimize intermolecular interactions, followed by a final explicit-solvent refinement that improves interface geometry and binding energies.

For our system, we incorporated interaction restraints derived from NMR epitope mapping experiments. Tau residues 371–386, corresponding to the epitope-forming region, were defined as active residues in HADDOCK to guide the docking toward biologically relevant antibody–tau complexes. The complementarity-determining regions (CDRs) of DC11 were identified from sequence using the AbRSA web server and were likewise defined as active residues of the antibody to ensure correct interface definition and to avoid non-specific tau binding. Representative structures from coarse-grained MD simulations of Tau321-391 were used as input ligand conformations [30]. We selected a representative structure from the most populated cluster (cluster 1), which corresponds to partially folded tau conformations featuring N-and C-terminal interactions, as well as the centroids of clusters 2 and 8, representing extended tau states. In the following text, these tau structures will be referred to as conformations 1, 2, and 8, respectively.

HADDOCK generated multiple Tau–antibody complex models, which were clustered by structural similarity and ranked by HADDOCK score (referred to as models in Table S1). Lower scores indicate more favorable predicted binding interactions. The best-ranked models along with their HADDOCK scores and additional parameters are summarized in the Supplementary table S1. For selecting clusters for further analysis, we also considered, in addition to the HADDOCK score, cluster size and RMSD values reflecting structural convergence within each cluster.

For each HADDOCK cluster, four models were selected, resulting in a total of 12 DC11– Tau321-391 complex structures for further analysis. Intermolecular contacts forming the binding interface and their frequencies across the resulting models were analyzed independently. Contacts present in ≥7 of 12 models were classified as key contacts, reflecting high reproducibility across the conformational ensemble. Each HADDOCK run yielded an ensemble of closely related binding poses rather than a single well-defined stable complex, as expected from the use of predefined active residues. However, despite the overall similarity in global docking orientation, the intermolecular contact patterns between antibody and tau varied across the models, indicating flexibility in the specific interaction network within the binding interface. As expected, contributions from the N-terminus were observed only in models derived from Tau321-391 conformation 1 (Fig. 7). In contrast, in models based on Tau321 conformations 2 and 8, the N-terminus is projected away from the antibody and does not participate in binding (Fig.S5).

**Figure 7.**
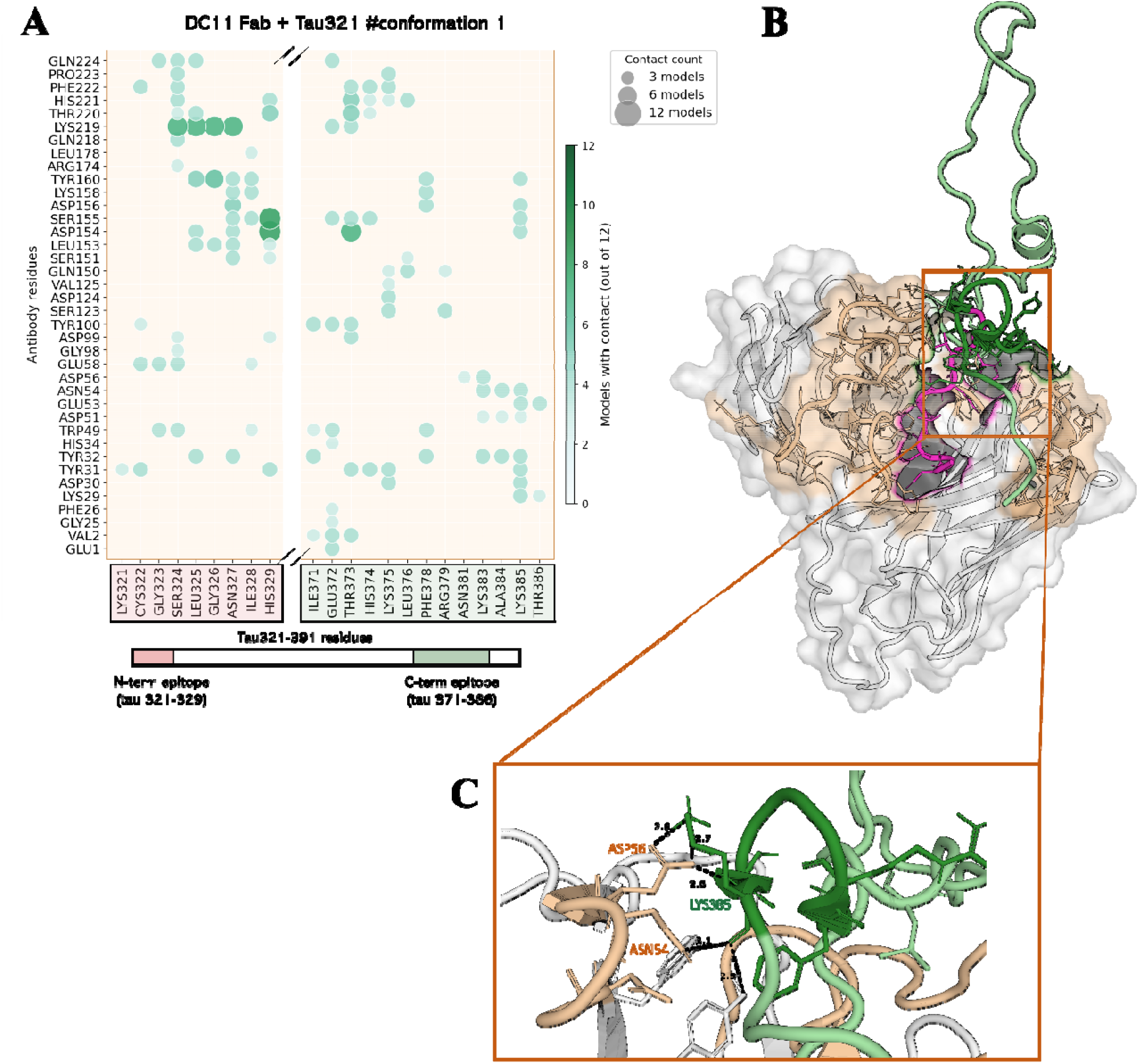
Antibody-Tau321-391 #conformation 1 interface analysis. (**A**) Quantification of antibody– tau interactions. Tau residues identified by NMR as constituting the primary DC11 epitope are highlighted in the green box. Additional N-terminal tau residues involved in complex formation are highlighted in the pink box. The two tau regions are separated by an axis break reflecting a sequence gap. **(B)** HADDOCK model of the DC11–Tau321 complex. The antibody is shown in surface representation, with CDRs colored light orange. Tau321 is shown as a cartoon representation in green, with N-terminal residues colored pink and NMR-identified epitope residues shown in dark green. Interacting residues are shown as sticks. (**C)** The local β-bridge structural motif formed within Tau321 C-terminal epitope residues.

Run 1 (DC11 + Tau321-391 conformation 1) differs from the other two docking runs due to the structural properties of the input tau conformation. This particular conformation, characterized by intramolecular contacts between the N-and C-termini, was selected to examine the involvement of N-terminal residues 321–325 in antibody binding. Intermolecular contact analysis revealed two potential epitope surfaces, consistent with the folded-back tau conformation observed in MD simulations [30]. Within the canonical C-terminal epitope, the most prominent interaction involved D154 (D31L) of DC11 and T373 of tau, whereas the remaining contacts in this region appeared more diffuse and variable across the models. In contrast, the N-terminal interaction map (Fig. 7) is dominated by contacts between DC11 residue K219 (K96L) and tau residues N327, G326, L325, and S324, which are present in 7 of 12 analyzed models. Together with consistent interactions between S155 (S32L) and D154 (D31L) of DC11 and H329 of tau (observed in 8 models), these constitute the highest-confidence contacts identified in this run.

A particularly interesting observation was the persistence of a β-bridge motif within the C-terminal epitope region spanning residues 378–383. This hairpin-like structural element was already present in the MD simulations, where it was found in the most populated cluster, accounting for 7.49% of the conformational ensemble (MD data published in Njemoga et al., 2026 [30,36]). Importantly, the motif was preserved after docking and remained engaged within the DC11 binding interface in models derived from Tau321-391 conformation 1.

At atomic resolution, the interface between the antibody and the C-terminal epitope revealed a dense network of interactions, centered primarily on tau residue K383. K383 formed a bidentate salt bridge with the two carboxylate oxygens of antibody residue D56 and also simultaneously engaged in a hydrogen bond with N54. This type of bidentate interaction is characteristic of particularly stable electrostatic contacts at protein–protein interfaces. The neighboring tau residue, A382, also contributed to binding via a backbone-mediated hydrogen bond to Y32 of the antibody. Taken together, the combination of salt bridges, hydrogen bonds, and backbone-mediated contacts within this short tau segment supports the presence of a specifically recognized structural motif. Moreover, the β-sheet propensity of the ^376^LTFRENA^383^ region of tau, as identified from several amyloid predictors, including TANGO, WALTZ or PASTA [30] suggests that DC11 may preferentially recognize and stabilize a locally ordered, hairpin-like conformation. Such a mechanism could help explain how the antibody discriminates between highly disordered tau states and conformations with increased aggregation propensity, while highlighting the A382–K385 region as a key structural determinant within the broader epitope.

Tau321 conformations 2 and 8 interacted with the DC11 paratope primarily through residues belonging to the defined C-terminal epitope in the remaining two HADDOCK runs. Similar to run 1, the interaction patterns were relatively sparse and heterogeneous, indicating a flexible binding interface rather than a single rigid binding pose, which is typical for complexes involving intrinsically disordered proteins (Fig. S5). Several interaction hotspots could nevertheless be identified from the run 2 contact map, mainly involving antibody residue K219 (K96L), which primarily interacts with tau residues E380 and L376. Similar to the lysine-centered interactions observed in run 1, tau residue K385 also interacted with antibody residues D56 and N54 (Fig. S5A).

K219 (K96L) again emerged as the dominant contact in run 3, mainly interacting with tau N381. In addition, run 3 revealed another cluster of recurrent interactions involving antibody residues Y31 and Y32 contacting tau residues K375, L376, and F378, together with contacts between W49 and F378. These interactions point to additional hydrophobic and aromatic packing contributions, particularly around F378, that may further stabilize the binding interface.

Overall, the docking analysis suggests that DC11–Tau321-391 recognition is driven primarily by electrostatic interactions, with lysine residues playing a central and recurring role on both sides of the binding interface. A common paratope hotspot centered on antibody residue K219 (K96L) was identified across all three binding modes, while an additional interaction hotspot involving tau residues A382–K385 coincided with the nascent β-hairpin motif observed in the N-terminal engagement mode.

## Discussion

The antibody DC11 is highly specific for misdisordered truncated tau proteins [7]. As such, structural insights into its binding tau epitope tau are of high importance, because this can inform on the pre-aggregation conformation of tau protein. Here, we report the high-resolution X-ray structure of DC11 antibody Fab and identify its tau binding epitope by ELISA, NMR and XL-MS methods. Building on the previously identified epitope, the truncated tau peptide tau321-391 [6], we sought to provide more accurate residue level data. All methods identified the C-terminal residue segment between residues 371-386 and also the N-terminal part 321-325 as part of the binding epitope. Of potential relevance is the latter sequence, previously shown to be important for the aggregation of truncated tau proteins, as tau321-391 was capable of amyloid formation without the need for heparin induction [30]. With molecular docking we have constructed a presumed model of the DC11-tau complex which fulfils the experimental constraints where both C and N terminal tau epitopes are involved in the interaction with antibody.

The structure of another anti-tau conformational oligomer specific antibody was published in its apo-form [37] as a scFv fragment of a previously crystallized rabbit antibody [38]. This antibody was shown to inhibit seeding of monomeric tau by brain extracts from AD patients and was crystallized as a monomer, dimer and a trimer of scFvs.

Inert and seed-competent tau monomers were previously identified in the conformational ensemble of tau protein [39]. The seed competent tau conformation has the aggregation prone sequences VQIINK and VQIVYK exposed

Crystal structures of the MN423 Fab fragment alone and in complex with its tau epitope revealed an identical antibody combining site configuration in both forms, suggesting that MN423 acts as a rigid structural mold that imposes a defined conformation on the disordered tau segment upon binding [40]. The fact that the antibody binding site is pre-formed even in the absence of the peptide implies that tau itself is capable of adopting this conformation spontaneously in the presence of MN423. In contrast, the binding of tau peptide to the antibodies Tau5 and DC8E8 is accompanied by conformational changes of the antibody combining site (PDB IDs 4TPR, 4TQE, 5MX3, 5MO3), where the antibody binding site seems to accommodate preformed small structural motifs (S-turn in the case of tau5 antibody) in tau protein [41]. As antibodies MN423 and DC11 potentiate in vitro tau aggregation, they should be able to imprint an aggregation prone conformation to the molecules of monomeric tau in solution, implying an induced fit binding mechanism.

The length of the antibody CDRH3 loop in mouse ranges from 4 to 28 with an average length of (11.5□±□1.9 aa residues), so the antibody DC11 features one of the shortest known CDRH3 lengths [42]. Its binding epitope on aggregated tau seems to be located at the interface between the structured core (ending upmost at residue E380 [43]) of the tau filament and its fuzzy coat.

## Funding

This research was funded by the EU NextGenerationEU through the Recovery and Resilience Plan for Slovakia under the project No. 09I03-03-V04-00623. This work was supported by research grants VEGA 2/0125/23 and 1/0825/25. This research was further funded by the European Union’s Horizon Europe program under the grant agreement No. 101087124.

CIISB, Instruct-CZ Centre of Instruct-ERIC EU consortium, funded by MEYS CR infrastructure project LM2023042 and European Regional Development Fund-Project “Innovation of Czech Infrastructure for Integrative Structural Biology” (No. CZ.02.01.01/00/23_015/0008175), is gratefully acknowledged for the financial support of the measurements at the Josef Dadok National NMR Centre. Initial NMR measurements were funded by iNext Discovery.

We gratefully acknowledge Dr. Ondrej Šedo and colleagues from CEITEC proteomics CF for LCMS/MS analyses.

