## Supplementary file for "Structure and epitope mapping of the conformational anti tau antibody DC11"

**Table S1.** Top-ranked HADDOCK clusters obtained from DC11-Tau321-391 docking selected for further analysis.

| **Docked Tau321 conformation from MD clusters** | **HADDOCK model** | **HADDOCK score** | **Number of structures** | **RMSD** |
| --- | --- | --- | --- | --- |
| #conformation 1 | cluster 8 | -100.44 | 6 | 0.80 |
|  | cluster 2 | -95.97 | 20 | 1.78 |
|  | cluster 1 | -90.52 | 29 | 0.97 |
| #conformation 2 | cluster 1 | -114.1 | 48 | 1.36 |
|  | cluster 2 | -105.6 | 27 | 1.64 |
|  | cluster 4 | -98.5 | 16 | 2.33 |
| #conformation 8 | cluster 3 | -117.3 | 18 | 1.21 |
|  | cluster 1 | -100.9 | 100 | 2.83 |
|  | cluster 5 | -93.7 | 8 | 2.46 |

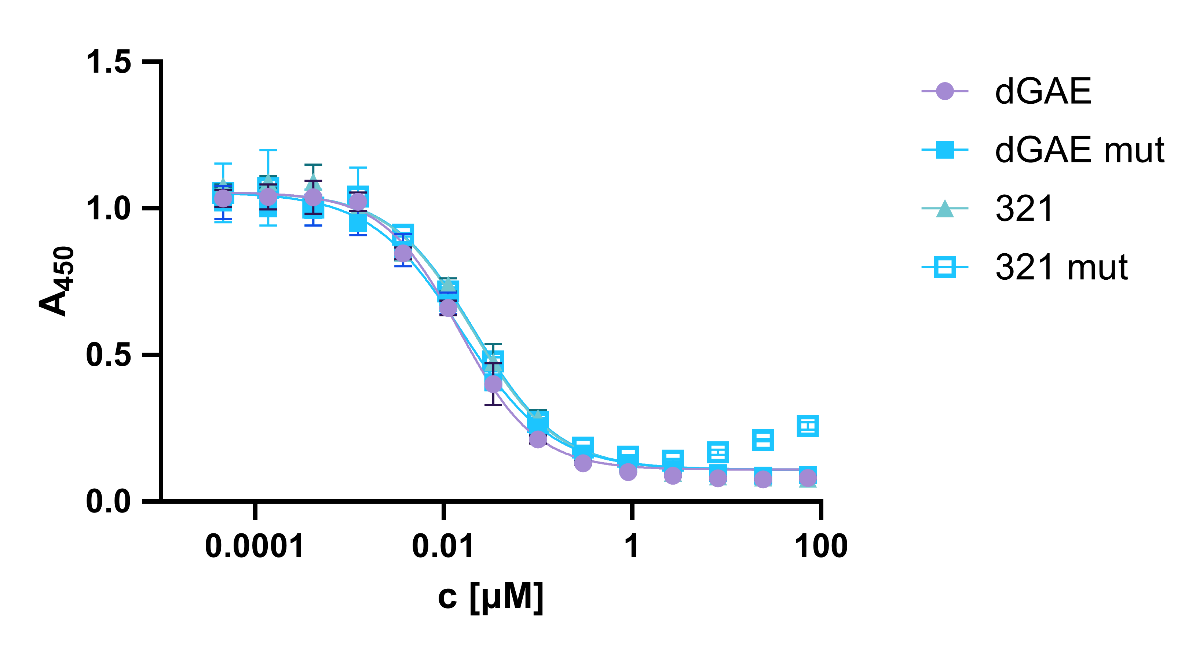

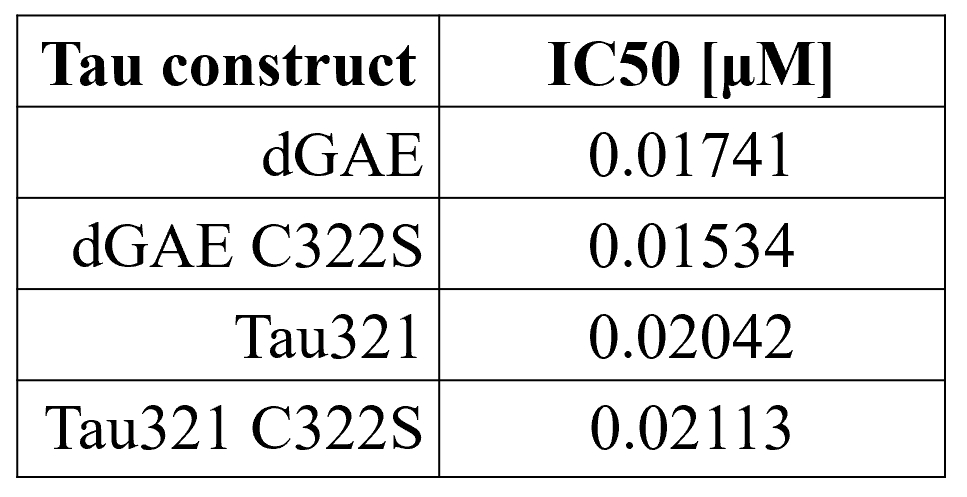

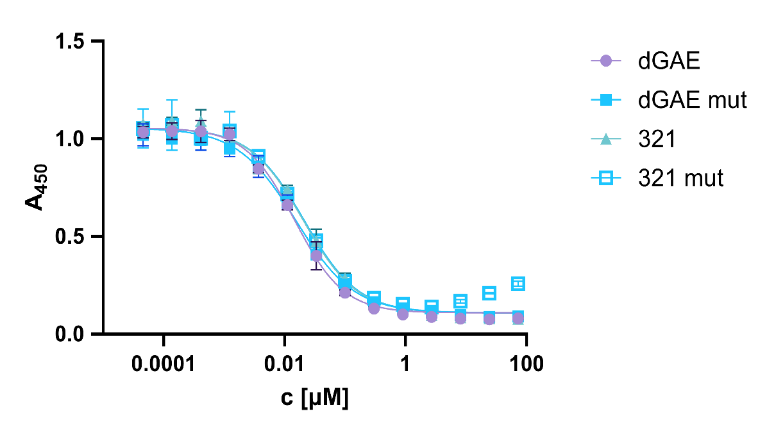

**Figure S1: Competitive ELISA of wild-type and C322S tau variants**. Increasing concentrations of soluble competitor proteins led to a concentration-dependent reduction in absorbance at 450 nm. Wild-type and C322S variants displayed highly similar inhibition curves, indicating that substitution of Cys322 has little effect on antibody recognition. Error bars represent SD from duplicates.

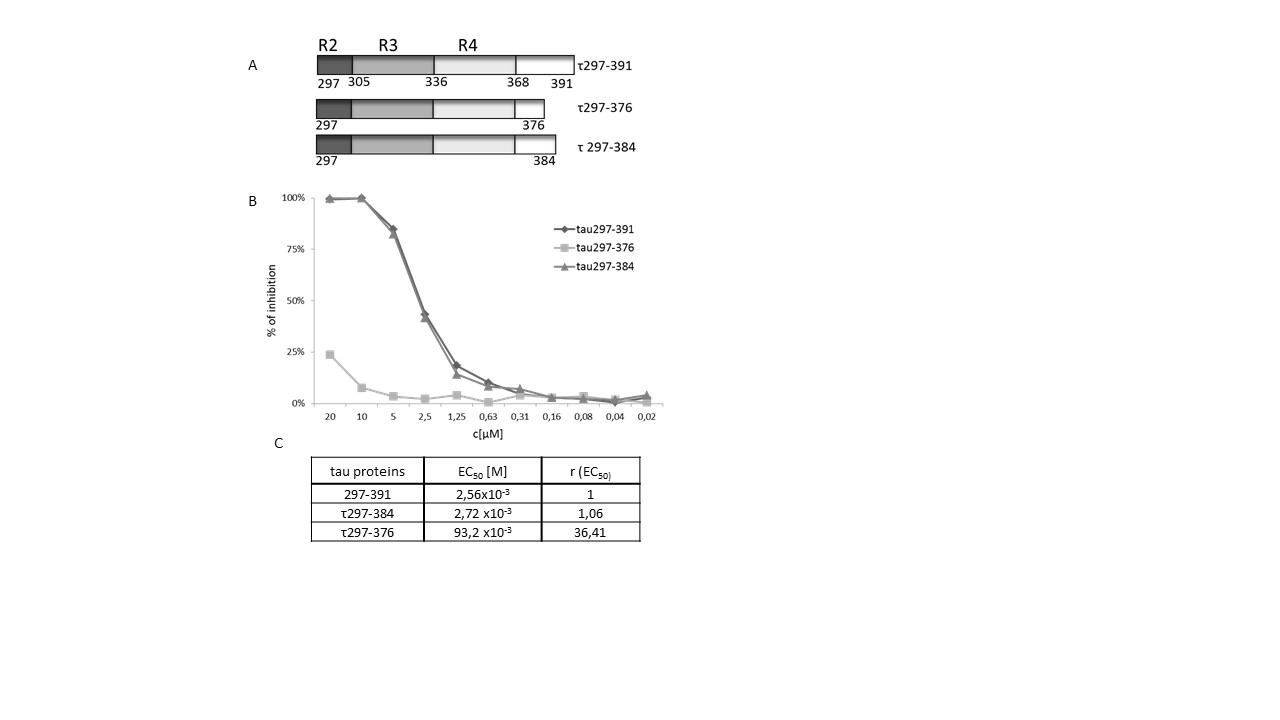

**Figure S2:** Identification of sequence contributing to the epitope of DC11 antibody located in C-terminal sequence 368-391. A) Structural diagram of C -terminal deletion mutants derived from τ297-391 used for immunoreactivity of DC11 in competitive ELISA. B) Figure shows competition of C-terminal deletion mutants of τ297-391 for binding with DC11. Individual competition curves reflect immunoreactivities of respective mutants. EC_50_ values were calculated for measure the affinity of DC11; ratio between EC_50_ of respective deletion mutant and EC_50_ of τ297-391 ( r ) indicates decrease in relative affinity of mutant (r-fold) in relation to full affinity of τ297-391 .

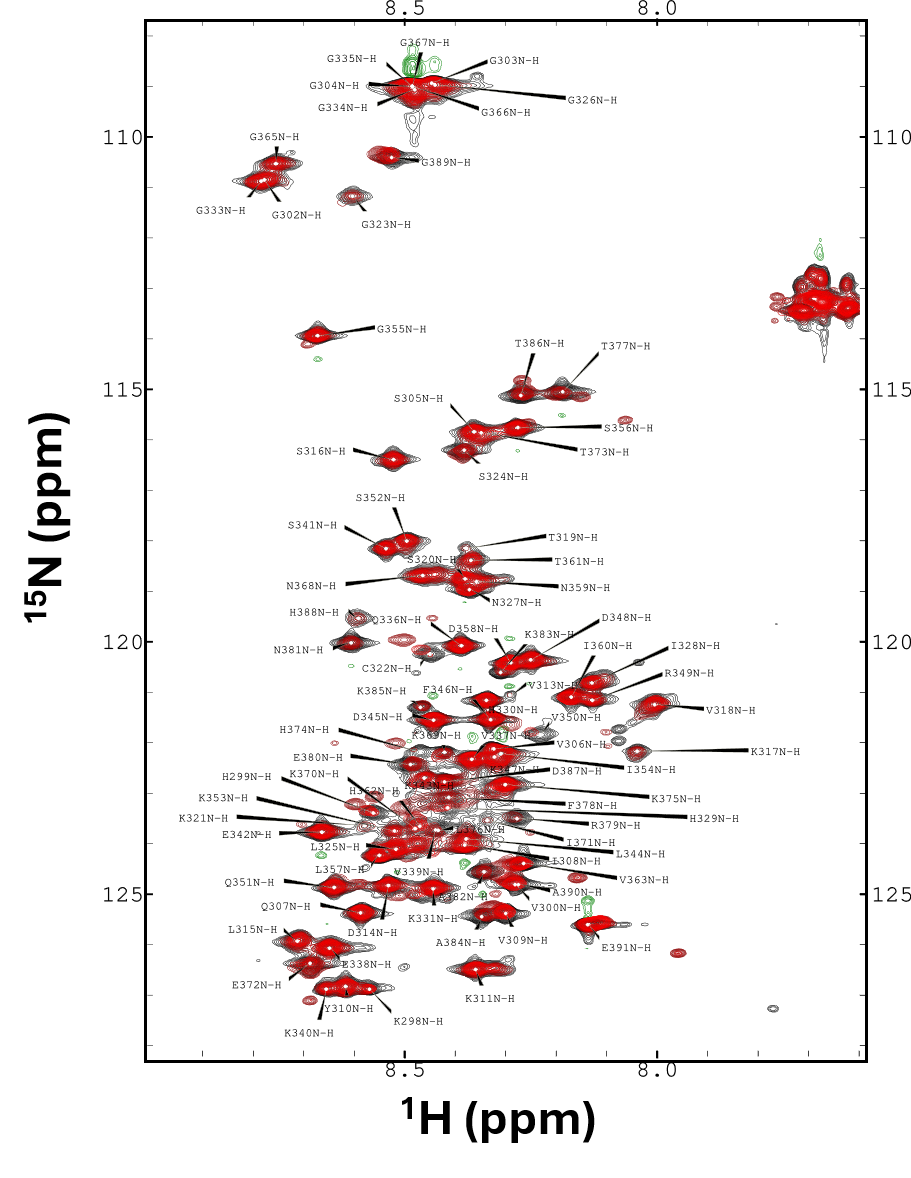

**Figure S3 - 2D HSQC NMR spectrum of [^13^C,^15^N]** **dGAE titrated with DC11 FAB.** Annotations show the assignment of amide backbone N-H correlations for all residues with the exception of N-terminal M296. Positive contours are coloured black to red with increasing DC11 FAB ratio (1:0 to 4:1, see methods for details) following the colour scheme in Fig. 6A.

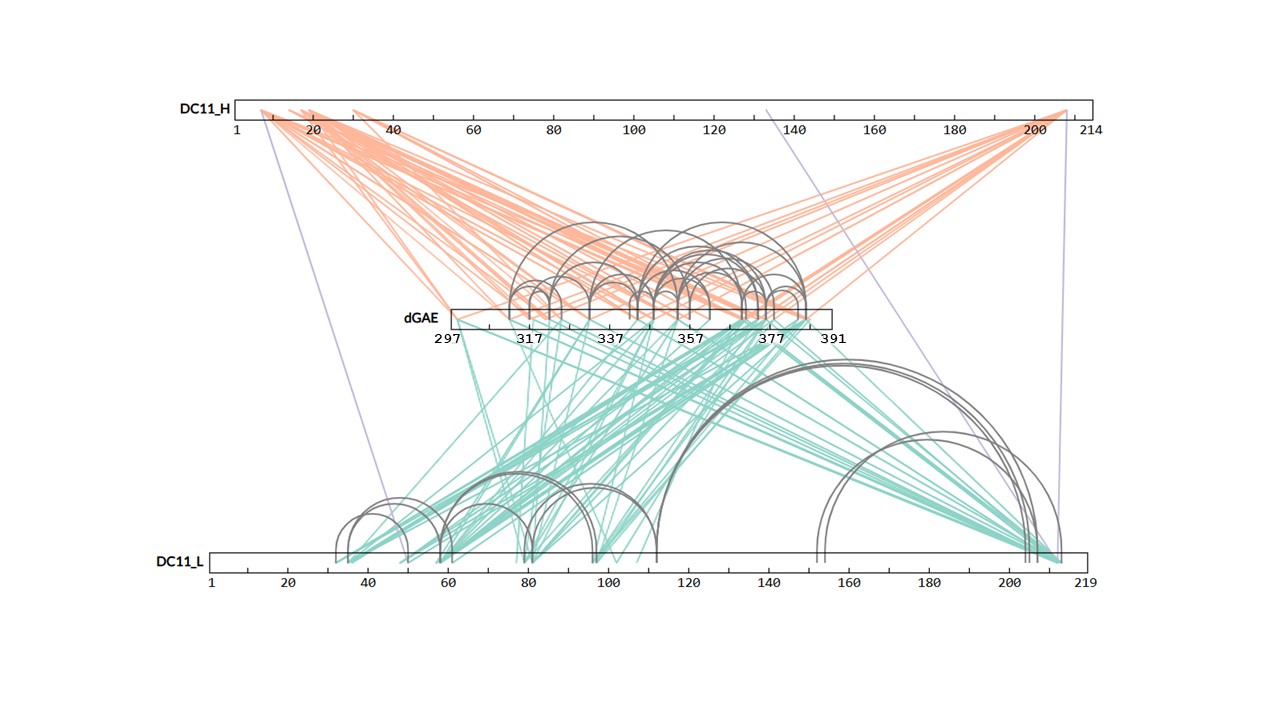

**Figure S4 - Unique crosslinks detected in XL-MS experiments.** Intramolecular crosslinks are shown between dGAE and the heavy chain (DC11_H, orange lines) or the light chain (DC11_L, green lines) of DC11 FAB, in addition to links between DC11 chains (blue lines) and intermolecular crosslinks (grey lines).

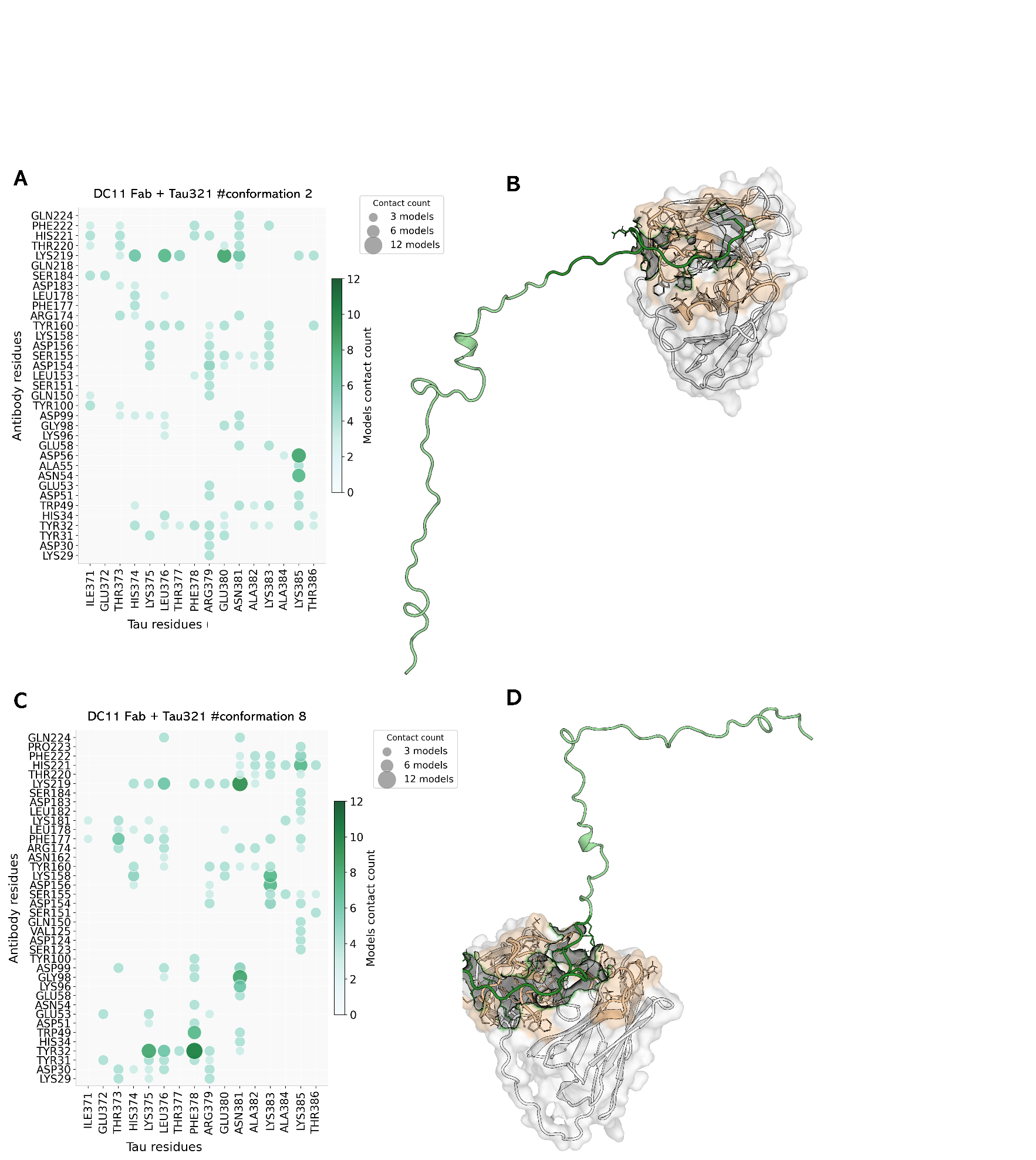

**Figure S5: DC11 Fab-Tau321-391 #conformations 2 and 8 interface analysis.** A, C: Quantification of antibody–tau interactions. B, D: HADDOCK models of the corresponding DC11–Tau321 complex. The antibody is shown in surface representation, with CDRs colored light orange. Tau321 is shown as a cartoon representation in green, with NMR-identified epitope residues shown in dark green. Interacting residues are shown as sticks.
